# *Acinetobacter baumannii* detected on mCCDA medium in a waste stabilisation pond

**DOI:** 10.1101/367128

**Authors:** Maxim Sheludchenko, Anna Padovan, Mohammad Katouli, Anne Roiko, Helen Stratton

## Abstract

*Acinetobacter baumannii* survives for prolonged periods under a wide range of environmental conditions. In a larger study investigating the efficacy of pathogen removal in a waste stabilization ponds (WSP), we cultivated microbes from wastewater samples on mCCDA agar containing selective and recommended supplements for the growth of *Campylobacter*. This bacterium is a recommended reference pathogen for the verification and validation of water recycling schemes in Australia and other parts of the world. A high number of colonies characteristic of *Campylobacter* grew on the selective media but this did not correlate with qPCR data. Using primers targeting the16S rRNA gene, and additional confirmatory tests such as detection of VS1, *omp*A, bla_OXA-51-like_, bla_OXA-23-like_ genes, we tested eight random colonies from eight samples (64 colonies in total) and identified them as *A. baumannii*. Wastewater grab samples taken three times over 6 months throughout the WSP system showed removal of *A. baumannii* in the WSP atrates similar to *E. coli*. In contrast, further intensive sampling from the inlet and the outlet of the WSP using a refrigerated auto-sampler showed that the number of *A. baumannii* in most sampling rounds did not differ significantly between the inlet and outlet of the WSP and that there was high variation between replicates at the outlet only. Resistance genes were detected in most *A. baumannii* isolated from the waste stabilisation pond and may potentially be a source of antibiotic resistance for environmental strains.

## Introduction

Waste stabilization pond (WSP) systems are used widely for the treatment of domestic wastewater in developing countries due to low cost and easy maintenance^1^. A typical WSP system consists of two lagoon types: facultative and maturation pond(s). Often, raw sewage goes directly into a 1.5-2 m deep facultative pond with a retention time of about 7-10 days, after which the wastewater flows into a baffled or non-baffled maturation pond with a longer retention time. As reported earlier, pathogenic strains such as *Salmonella enterica* and *Campylobacter jejuni* could be removed in a facultative pond and may not be present any longer in the maturation pond effluent^2^. Nevertheless, it remains interesting how pathogenic *Campylobacter* spp. survive in the environment, as the bacterium is a foodborne pathogen and is also a major cause of water-borne diarrhoeal outbreaks^3^. The Australian water recycling guidelines (AWRG) recommend monitoring *Campylobacter* as a reference bacterial pathogen together with faecal indicators and other reference pathogens such as adenovirus and oocysts of *Cryptosporidium*^4^ However, routine testing for *Campylobacter* spp. in wastewaters using culture-based methods often yields biased results due to the growth of other species (Judy Blackbearrd, Melbourne Water, Australia; personal communication). Selective media such as Bolton, Preston and mCCDA agars for the detection of *Campylobacter* spp. have been used in many studies involving food–borne diarrheal outbreaks^5-7^ However, the use of these media and their specificity for detection of *C. jejuni* in environmental samples has been questioned ^8-10^. It is known that other bacterial species such as *E. coli, Proteus* spp., *Acinetobacter* spp. ^11^ as well as *Helicobacter* spp., *Arcobacter* spp., *Sutterella wadswortheness* and *Bordetella pertussis* ^12^ produce characteristic grey colonies of *Campylobacter* on these selective media and can be counted as false positives.

The present study was part of a larger investigation studying the degree of pathogen removal within WSPs in Australia. Using a combination of culture-based and molecular techniques to detect the presence of *Campylobacter* spp. in a WSP, we observed a high number of colonies with characteristics of *Campylobacter* spp. on mCCDA agar plates that did not corresponded with the results of qPCR on the same samples. We employed 16S rRNA sequencing together with species-specific primers to characterise a high number of these colonies. The results confirmed that these colonies were mainly *Acinetobacter baumannii*. We then performed an intensive sampling from the inlet and the outlet of this WSP using both grab sampling and auto-samplers and tested for the presence of *A. baumannii* using the above methods. The implications of collecting samples using auto-samplers and utilising the mCCDA medium for the identification of *C. jejuni* for the enumeration of pathogenic bacteria in the effluent of waste stabilization ponds is discussed, particularly in light of long term monitoring of *A. baumannii*.

## Results and Discussion

During a study investigating pathogen removal in waste stabilisation ponds, we noticed a lack of correlation between the number of presumed *Campylobacter* spp. based on mCCDA agar (supplemented with polymyxin B to reduce growth of Gram-negative bacteria) and qPCR data of the same samples. We suspected that the mCCDA agar plates were supporting the growth of non-campylobacter species and sequencing of the 16s rRNA gene of colonies isolated from the mCDDA plates showed that they were in fact *A. baumannii*.

Monthly replicate grab samples were collected from the inlet and six sites throughout the baffled WSP pond, as well as from constructed wetlands (CW) and reed beds (RB) that were a part of the treatment chain following the WSP. Analysis of these samples demonstrated a reduction in the number of *A. baumannii* as determined by membrane filtration on mCCDA agar and was strongly correlated with the faecal indicator, *E. coli* during all observations (Figure 1). To monitor pond performance, we also counted the number of *E. coli*, a conventional faecal indicator bacterium, in all samples. The major reduction of *A. baumannii* occurred at the beginning of the WSP system i.e. at sites H1 to H3 and where the effluent entered the RB after passing through CW, as described in the methods. The initial numbers of *A. baumannii* in influent samples were 2 logs lower than *E. coli*, though the removal rates remained the same, suggesting that *E. coli* can be used as an acceptable indicator for monitoring changes of *Acinetobacter* in such maturation ponds.

**Figure 1.**
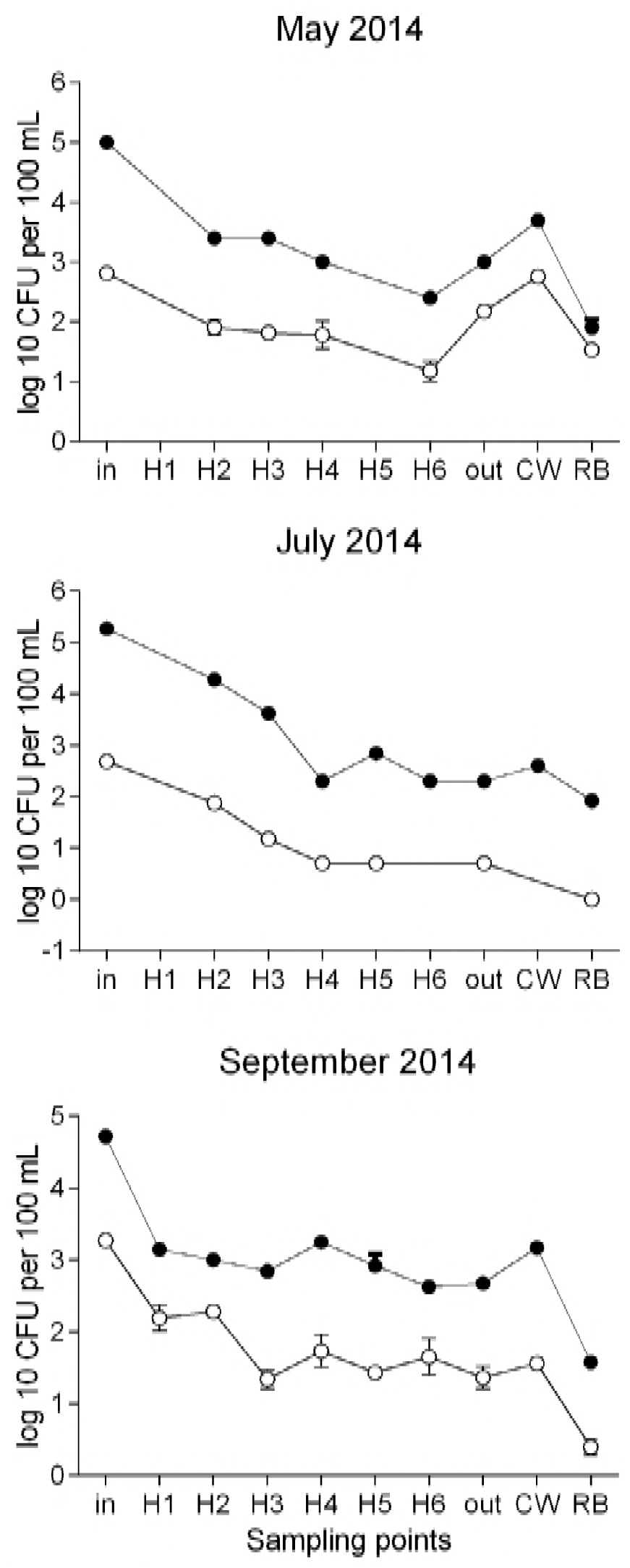
Concentrations of *A. baumannii* (○) and *E. coli* (●) in the maturation pond at the inlet, at multiple locations (H1-H6) between baffles, at the outlet and effluent from constructed wetlands (CW) and reed beds (RB). Bars indicate standard deviation.

Using the auto-sampler, we sampled wastewater intensively from the inlet and outlet of the WSP throughout February and March 2015 to monitor faecal indicators and pathogens over a long-time period. Most pond monitoring is done monthly but we wanted to determine whether bacterial concentrations and their removal rates changed over time. The number of *E. coli* in almost all samples collected from the inlet were between 1-2 logs higher than *A. baumannii* (Figure 2A) as was observed during grab sampling in 2014. *E. coli* numbers varied from 3.45 to 6.31 log in the inlet and from 1.15 to 3.92 log at the outlet with a consistent 2-3 log reduction in *E. coli* numbers observed throughout the sampling (Figure 2B). In contrast to the observations made in 2014 using grab samples, the reduction of *A. baumannii* varied in the effluent samples and the concentration of *A. baumannii* was mostly lower than that of *E. coli* (Figure 2B). During the first 48 hrs sampling in February (3 samples per day) we observed a 1-2 log reduction in the number of *A. baumannii*. However, from 10 February 2015 (the third round of sampling) onwards, the number of *A. baumannii* were higher (1-2 log) than *E. coli* despite their lower initial numbers at the inlet. In addition, a large variation in the numbers of *A. baumannii* was observed between duplicate samples from the effluent, with higher concentrations observed in the first duplicate and lower concentrations in the second duplicate. This was not observed for *E. coli*. One possible explanation for this higher variation between duplicate samples is that pond water at the outlet site had to pass through 7 meters of tubing before it reached the auto-sampler and this may have contributed to bacterial growth inside the tubing. The auto-sampler has a backwashing function to avoid collecting remnant water, however, it may not have worked sufficiently to eliminate these bacteria inside the tubing, causing higher numbers in the first sample. We postulate that the higher numbers of *A. baumannii* in outlet samples compared to the inlet could also be due to the biofilm formed by these bacteria inside the tubing. *Acinetobacter* spp. can form biofilms in different environments^19-21^. Thus, the use of auto-samplers with long tubing may not be suitable for continuous monitoring of bacterial pathogens which can form biofilms.

**Figure 2.**
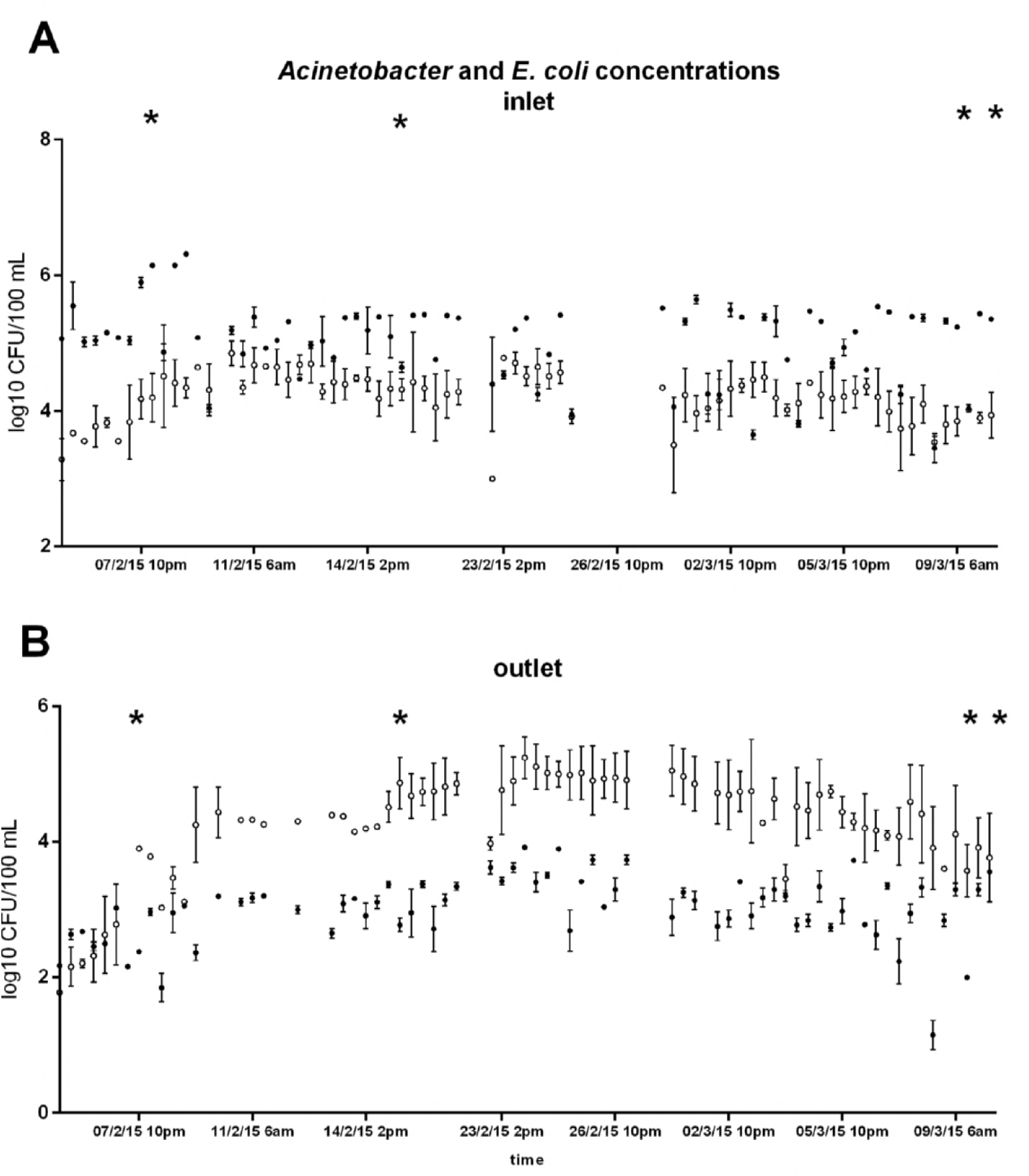
*A. baumannii* (○) and *E. coli* (●) concentrations (log10 CFU per 100 ml) in the inlet (A) and outlet (B). Samples were taken at 8 hourly intervals with refrigerated auto-samplers, at 6 am, 2 pm and 10 pm. Bars indicate standard deviation. *Confirmatory screening by real-time PCR was undertaken for colonies from these samples and results are presented in Figure3.

To rule out the possibility that some of the isolates on the mCCDA plates could have been *C. jejuni* or other species such as *C. ureolyticus*, the second most common species of *Campylobacter* after *C. jejuni* in faecal specimens of patients ^22^ and some animals ^23^, we tested 539 isolates from mCCDA plates over four sampling events. The results showed that none of the colonies tested were positive for the *C. jejuni* VS1 specific marker or any of the other 14 molecular markers specific for other species of *Campylobacter* (see Supplementary Materials, Table S1).

*Acinetobacter* spp. and in particular *A. baumanii* have been reported in wastewater effluent ^24,25^ with antibiotic resistance shown to increase in these bacteria due to the selective pressure of the environment ^26^. In the WSP studied here, we also identified a range of pharmaceuticals including antibiotics (see Supplemental Material Table S2). Therefore, we investigated antibiotic resistance of *A. baumannii* isolates from mCCDA plates by PCR, which was also another confirmation step for the identification of *A. baumanii*. Between 7^th^ of February and 10^th^ of March of 2015, an additional five rounds of grab sampling were performed at the inlet and outlet of the WSP. In all, 270 isolates from the inlet and 216 isolates from the outlet were tested for the presence of the *A. baumannii* specific antibiotic resistance markers *omp*A and two beta-lactamase genes i.e. *oxa*-51 and *oxa*-23. The results showed that 96% of the isolates were positive for the *omp*A marker (Figure 3). Shotgun metagenomics analysis also confirmed the presence of antibiotic resistant sequences of *A. baumanii* in DNA extracts from the same water samples.

**Figure 3.**
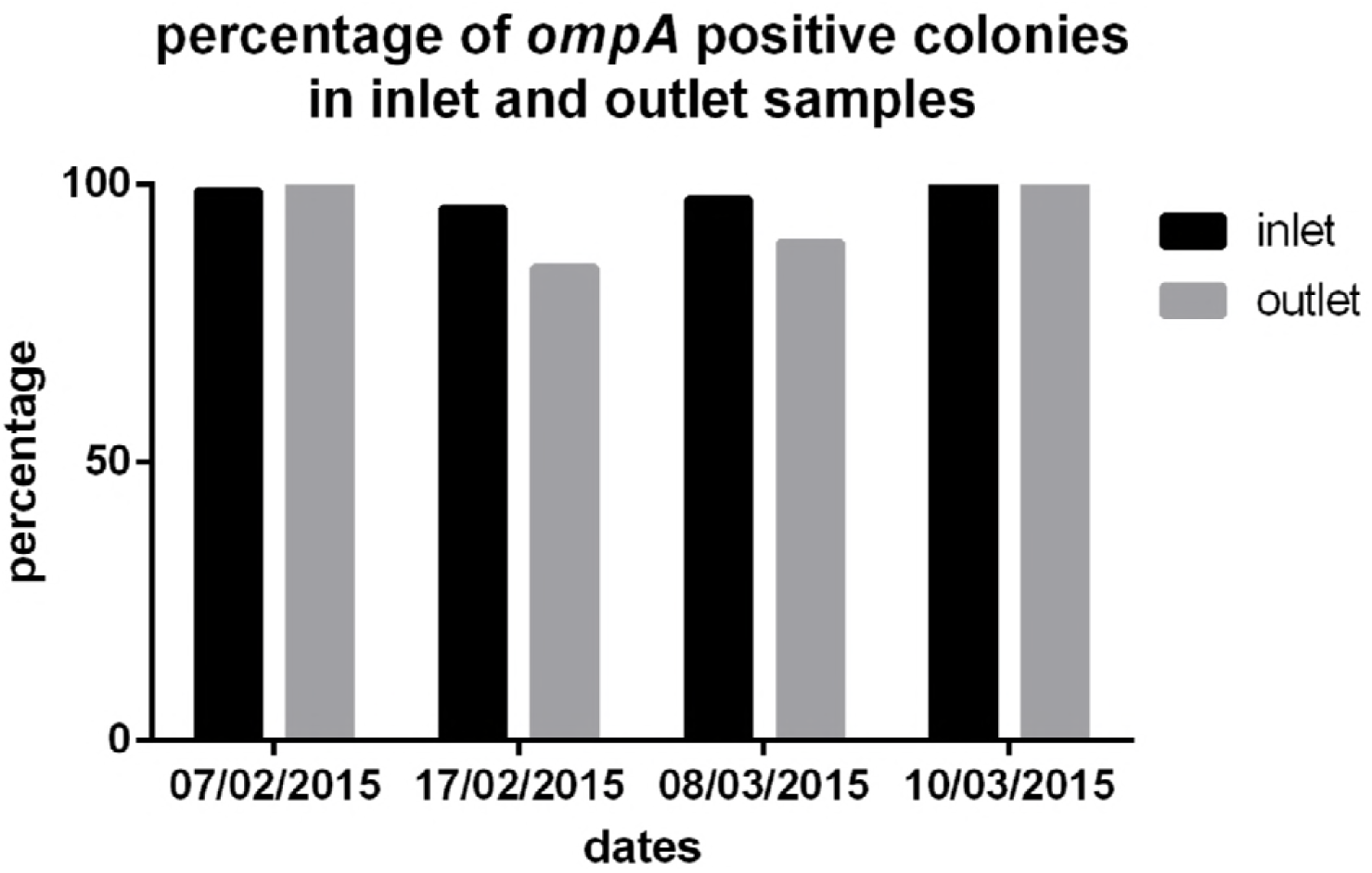
Percentage of *omp*A positive colonies out of the total number of isolates tested (Table S1) on different occasions during 30 days of monitoring using auto-samplers. DNA was extracted from colonies isolated on mCCDA medium originating from wastewater collected at the inlet and outlet, and tested by PCR.

A specific characteristic of *A. baumanii* strains is their susceptibility to carbapenemase antibiotics ^17^. In our study, two of seven colonies taken from the same plate were positive for *bla*A_oxa-51_ and negative for *bla*A_oxa-23_. Thus, we also confirmed the presence of *A. baumanii* in effluent wastewater with carbapenemase resistance. Recently, the World Health Organization (WHO) raised concerns about the presence of antibiotics in wastewater and the consequences of transfer of antibiotic resistance genes to environmental bacteria upon their exposure^27^. For this reason, *A. baumanii* maybe an important candidate for monitoring bacterial pathogens in WSPs.

To the best of our knowledge, this is the first report on the presence of antibiotic resistant strains of *A. baumanii* in a WSP system. The presence of a range of pharmaceuticals in the wastewater effluent from the studied maturation pond (see Supplemental Material Table S2) may contribute to the presence of antibiotic resistant strains of *Acinetobacter*. *A. baumannii*, despite being a weak pathogen compared to *Pseudomonas aeruginosa*, may play a significant role in spreading broad-spectrum resistance genes to other Gram-negative organisms ^28^, especially under environmental conditions of waste stabilization ponds. In Australia’s northern tropics, these bacteria are responsible for major cases of community-acquired pneumonia during the wet season ^29^. The presence of clinically important strains of *Acinetobacter* in the environment is not known but few authors have reported antibiotic resistant bla_OXA-23_ *A. baumannii* in hospital wastewaters ^30^, in filaments of active sludge of wastewaters ^31^ and in paleosol contaminated with waste leakage ^32^. *A. baumanii* can also be very persistent in the environment, requiring special treatment for inactivation ^33^.

In conclusion, we found that *A. baumanii* could grow on selective mCCDA plates supplemented with polymyxin B forming colonies characteristic of *Campylobacter*. Therefore, the use of this medium for the detection and enumeration of *Campylobacter* spp.in wastewater or recycled water should be always accompanied by additional confirmatory tests. We also found that *A. baumanii* is highly abundant in the studied maturation pond in subtropical Australia and some of these strains might carry antibiotic resistance genes. Finally, the use of auto-samplers may not be suitable for continuous monitoring of *A. baumanii* in wastewater systems due to their ability to form biofilms.

## Material and Methods

### 2.1. Site description and sample collection

The WSP is located in subtropical South-East Queensland within a sub-urban community of 1500 inhabitants. Raw sewage with brief mechanical treatment is piped into a primary 65 × 65 × 1.5 m facultative pond and subsequently piped to a 30 × 60 × 1.2 m baffled maturation pond with 12-20 days of retention time for this pond. Two 30 × 60 m constructed wetlands (CW) and three 15 × 20 m reed beds (RB) aid to polish (further treat) the effluent of maturation pond.

Grab sampling was initially carried out on May, July and September 2014. Samples were collected in 500 mL bottles directly from the inlet pipe of the influent (i.e. Inlet), at 30 cm depth of the effluent pond at two and a half meters from the inlet site (H1), the edge of five baffles (H2 to H6) as well as within 2 m of the outlet pipe (i.e. outlet). The second round of sampling was done in 2015 where two portable Avalanche refrigerated auto-samplers (Teledyne ISCO, US) were used to collect duplicate samples of pond water (900 mL each) at the inlet and outlet of the pond. Samples were collected by the auto-sampler every 8 hours during February-March 2015: 6:00 am, 2:00 pm and 10:00 pm, over 30 calendar days. All samples were sent to the laboratory on ice every second day to ensure analyses were carried out within 24-48 hours. At each sampling event, the averaged values of duplicate bacterial concentrations were used for data analysis, with missing values disregarded. Overall, there were 73 inlet data points and 71 outlet data points for enumeration of *E. coli* (colonies on mTEC agar) and *A. baumannii* (colonies on mCCDA plates).

### Culture-dependent enumeration by direct plating

Individual wastewater samples were mixed thoroughly, serially diluted in sterile distilled water and filtered through 0.45 μm sterile mixed cellulose ester membrane filters (Advantec, Tokyo, Japan) using a CombiSart® manifold filtration unit (Sartorius, Gottingen, Germany). For enumeration of *E. coli*, as an indicator bacterium, filters were placed on modified mTEC agar (BD, Sparks, AR, USA) and incubated at 35°C for 2 hours followed by incubation at 44.5°C for 24 hours. Magenta colonies were counted as *E. coli*.

For enumeration of *Campylobacter* spp. and *Acinetobacter* spp., filters were placed on blood-free modified charcoal cefoperazone dexycholate agar (mCCDA, PO5091A, Oxoid, Basingstoke, UK) with antibiotic supplement (SR0155, Oxoid) as specified by the manufacturer. The antibiotic polymyxin B (SR0099, Oxoid) was also added to improve the manufacturer’s protocol as recommended by Chon *et al*.^11^. Selective chromogenic-like Brilliant CampyCount agar (PO1185, Oxoid) plates were obtained for a trial with wastewaters samples. The selective mCCDA plates were combined with an enrichment step using Preston broth (Nutrient broth N° 2, CM0067, Oxoid) which was supplemented with 5% lysed horse blood (SR0048, Oxoid) and antibiotic (SR0204 and SR0232E, Oxoid). Ten to fifty mL of influent and effluent were filtered through 0.45 μm sterile mixed cellulose ester membrane filters (Advantec) using a CombiSart® manifold filtration unit (Sartorius). The membrane was either directly placed onto selective plates or aseptically folded into 45 mL enrichment broth in 50 mL sterile plastic tubes. The tubes were tightly closed and left for 48 hours at 42°C. After this incubation step, 200 μL of the content was streaked onto mCCDA plates. Plates were incubated at 43°C for 48 hours under microaerofilic conditions using Campygen system (CN0025, Oxoid). All grey colonies were counted and eight of them were selected for further analyses.

### Identification of isolates by 16S rDNA amplification

Universal primers 27F and U1492R ^13^ were used to produce 16SrDNA amplicons in a PCR assay containing 1 × DreamTaq™ Master Mix (ThermoFisher Scientific, Sydney, Australia), 500 nM forward and reverse primer, 5 μL of DNA template in a total volume of 50 μL. Amplification reaction was performed in a C1000 thermal cycler (BioRad, Hercules, USA) with the following conditions: 95°C for 10 min, followed by 25 cycles of 95°C for 10 s, 50°C for 5 s and 72°C for 1 min with a final elongation step at 72°C for 10 min. PCR products were visualized on a 1% agarose gel stained with ethidium bromide and tubes containing PCR products of the correct size were purified using the QIAquick PCR Purification kit (Qiagen, Dusseldorf, Germany). Purified amplicons were directly sequenced (AGRF, Brisbane, Australia). The resulting 16SrDNA sequences were queried using the BlastN algorithm in GenBank for strain identification. Sequences were deposited in GenBank with accession numbers KU16092 and KU161093.

### Gene specific real-time PCR screening

Sixty four randomly selected colonies were re-grown on mCCDA, transferred to 96 well plates containing 100 μL of DNAse/RNAse free water using sterile plastic bacterial loops (Sarstdat, Germany) and mixed vigorously to disrupt the bacterial cells. The cells were boiled in a water bath for 10 min as described ^14^ and centrifuged at 4000 ×g for 10 min to pellet cell debris. The supernatant (70 μL) was transferred to new plates with 70 μL of DNAse/RNAse free water and stored at −20°C until further use.

Each qPCR reaction mixture contained 12.5 μL of 2× SYBR Select Master Mix for CFX (Life Technologies, USA), 400 nM each primer (Macrogen, Seoul, South Korea), and 5 μL of DNA template made to 25 μL with DNAse-RNAse free water. VS1 primer pairs for *C. jejuni* enumeration ^15^ and *cpn*60 primer sets for the identification of other strains of *Campylobacter* spp.^16^ were used. The *omp*A gene was targeted for the identification of *Acinetobacter* spp. and *A. baumanii* specific antibiotic resistance genes, bla_oxa-51_*bla*_oxa-23_, were used as previously described ^17, 18^. Real-time PCR reactions were placed into a CFX96 thermal cycler (Bio-Rad) with an initial polymerase activation of 95°C for 10 min, followed by 40 cycles of 95°C for 10 s, 60°C for 1 min with a final melting step of 60°C to 95°C, increasing 1°C at each cycle. Reactions assays for unknown samples were run in duplicate.

Positive controls used for PCR screening were *C. jejuni* NCTC 11168 and *A. baumanii* ATCC 19606. Bacterial cells were harvested by centrifugation at 3000 ×g for 10 min, washed with 1 × PBS and DNA was extracted using the FastDNA® SPIN kit for soil (MP Biomedicals, Santa Ana, CA).

### Shotgun sequencing

Thirty to 50 mL of wastewater collected in September 2014 was filtered onto 0.45 μm sterile mixed cellulose ester membrane filters membranes and DNA extracted using the FastDNA SPIN® kit for soil (MP Biomedicals). Extracts were outsourced for shot-gun sequencing with MiSeq 2×250bp (MR DNA, Shallowater, Texas). DNA libraries were prepared using Nextera DNA sample prep kits (Illumina, US). Sequences were uploaded and analysed with the MGRAST metagenome analysis server. Reads with size of 1,419,250 – 1,798,380 were given designation numbers 4666554.3-4666561.3 on the server.

## Acknowledgements

This study was supported by the Australian Water Recycling Centre of Excellence and the Queensland Science Fund Support. The assistance of the operational staff from Queensland Urban Utilities is greatly appreciated.

## Transparency Declarations

None to declare

## Conflict of Interest

The authors declare that there is no conflict of interest with the organization that sponsored this research and publications arising from this research.

